# N-terminal region of PetD is essential for cytochrome *b6f* function and controls STT7 kinase activity via STT7-dependent feedback loop phosphorylation

**DOI:** 10.1101/2025.02.21.639470

**Authors:** Afifa Zaeem, Yuval Milrad, Simon Bütfering, Carolyne Stoffel, Adrien Burlacot, Martin Scholz, Felix Buchert, Michael Hippler

## Abstract

The cytochrome *b_6_f* complex (*b_6_f*) links photosystem I (PSI) and photosystem II (PSII) in the photosynthetic electron transport chain and is distributed across appressed and non-appressed thylakoid membranes. The *b_6_f* also activates the state-transition 7 kinase (STT7), which phosphorylates light-harvesting complex proteins, triggering their migration to enable energy redistribution between PSII and PSI. STT7-dependent phosphorylation has also been observed at T4 in the N-terminal domain of the *b_6_f* subunit-IV (PetD), though its functional significance has remained unclear. To investigate its role, we generated several chloroplast mutants. The phosphomimic mutation PetD T4E – but not T4A – inhibited STT7 kinase activity, as indicated by the absence of STT7-dependent phosphorylation and a State 1-locked phenotype. This reveals a novel feedback mechanism regulating STT7-dependent phosphorylation. Deletion of the first five N-terminal amino acids similarly inhibited STT7 activity and additionally disrupted electron transfer, underscoring a crucial role of the PetD N-terminus in *b_6_f* function.

## Main

Photosynthesis captures light through photosystem (PS) I and PSII along with light-harvesting complexes (LHCI and LHCII). Light energy drives water splitting at PSII and ultimately reduces ferredoxin (FDX) through PSI. In linear electron flow (LEF), reduced FDX transfers electrons to FDX-NADP(H) oxidoreductase (FNR), producing NADPH for CO_2_ fixation and biomass production. In addition, electron transport deposits H^+^ in the lumen and separates charges across the thylakoid membrane, thereby generating both a proton gradient (ΔpH) and membrane potential (ΔΨ), which collectively drive ATP synthesis. Electron flow between the photosystems is mediated by the cytochrome *b_6_f* complex (*b_6_f*; reviewed in ^1^), which shuttles electrons from plastoquinol (PQH_2_) in the plastoquinone (PQ) pool to plastocyanin. Besides small peripheral subunits, the *b_6_f* core comprises cytochrome-*f* (PetA), cytochrome-*b_6_* (PetB), Rieske iron-sulfur protein (ISP, PETC), and subunit-IV (PetD). Electron flow through the high-potential chain, involving PETC and PetA, is functionally linked to redox reactions in the low-potential chain of PetB, including the heme *c*_i_ cofactor at the Qi-site. The Qi-site serves as the stromal binding pocket for PQ, whereas PQH_2_ binds at the lumenal Qo-site. By translocating protons into the lumen through reversible PQ protonation at the Qi and Qo sites, the ΔpH generated by *b_6_f* activity substantially contributes to ATP production required for CO_2_ fixation. Moreover, the *c*_i_ heme within the Qi-site is relevant for cyclic electron flow (CEF) and the *b_6_f* complex may function as a FDX-PQ reductase (FQR) in CEF ^2,3^.

The lumen acidification stemming from photosynthetic electron flows is important for efficient photoprotection under conditions of excess excitation energy: it establishes “photosynthetic control” at the Qo-site where PQH_2_ oxidation (and therefore electron flow) slows down at acidic pH ^4,5^, and it decreases photochemistry by stimulating controlled heat dissipation via non-photochemical quenching (NPQ) ^6–8^. Moreover, the *b_6_f* is linked to state transitions – another photoprotective mechanism to cope imbalanced excitation ^9^. During this process, LHCII migrates between PSII and PSI upon its phosphorylation by the thylakoid-bound STT7 kinase ^10,11^. STT7 is activated by a reduced PQ pool while specific phosphatases act as STT7 antagonists ^12^. The activity of the STT7 kinase is regulated by its transmembrane domain, with a disulfide bond between two lumenal cysteine residues being essential for its function ^13^. Active discussions and research on the kinase redox regulation are ongoing ^14–16^. Recently it was shown that state transitions involve PetD residues N122, Y124, and R125 in the stromal “fg Loop”, linking helices F and G of this *b_6_f* subunit ^17^. Moreover, the C-terminus of the *b_6_f* subunit PetB (PetB^C-term^) interacts with the fg Loop in PetD, thereby facilitating STT7 activation ^18^. Mechanistically, the fg Loop and PetB^C-term^ allow for binding of STT7 to *b_6_f* which enhances kinase autophosphorylation. In this report we focus on another stromal element of *b_6_f* in the model green microalgae *Chlamydomonas reinhardtii* (see Extended data Fig. 1): The N-terminus in PetD, where T4 undergoes phosphorylation in an STT7-dependent manner ^19^ (see also Extended data Fig. 2).

PetD T4 phosphorylation has been described by several groups ^19,20^ and to investigate the functional significance of STT7-dependent phosphorylation at PetD T4, we created PetD site-directed mutants T4A and T4E via chloroplast transformation to mimic constitutive absence and presence of phosphorylation, respectively. Moreover, the function of the five N-terminal residues was addressed in a truncation mutant (ΔN) whereas the wild-type PetD (WT) was used as control. The *petD* recipient strain carried a six-histidine tag at the C-terminus of PetA ^21^ for optional *b_6_f* purification. Spot test on agar plates, requiring photoautotrophic growth, revealed that mutant strains T4A and T4E performed similar to WT under normal light (40 μmol photons m^-2^ s^-1^, Extended data Fig. 3a). In moderate high light (200 μmol photons m^-2^ s^-1^; Extended data Fig. 3a), T4E revealed a severe growth phenotype in comparison to T4A and WT. In contrast, the ΔN strain showed a growth deficit at both light intensities compared to the other strains. Proteomic analyses showed no significant differences in the levels of PetD and other core *b_6_f* subunits between WT, T4A, and T4E. In contrast, most of these subunits were slightly depleted in ΔN, and the CEF effector protein PETO ^22^ was significantly downregulated (Extended data Fig. 3b). LHCSR1 – required for pH-dependent NPQ ^23^ – was markedly upregulated in T4E relative to T4A (Extended data Fig. 4). Moreover, STT7 peptide abundance was slightly lower in T4E compared to the similar levels observed in the other PetD variants (Extended data Fig. 5a). Given the observed growth and protein abundance phenotypes, key photoprotection pathways associated with *b_6_f* function might be altered in the mutants. To this end, we first present data investigating state transitions as they require the physical presence of *b_6_f* ^24^, and then performed electron transfer measurements to directly assess *b_6_f* activity in the mutants.

To explore if state transitions are affected in the mutants, the antenna redistribution between PSI and PSII was measured using 77 K fluorescence emission spectroscopy ^25^ after cells were shifted to either State 1 or State 2 inducing conditions (Fig. 1a–1d). Thereby, an increase in PSI-associated antennae size under State 2 versus State 1 conditions was observed for WT and T4A (Figs. 1a, 1b). In contrast, T4E and ΔN strains displayed no differences in their 77 K fluorescence emission spectra, indicating they were unable to perform state transitions (Figs. 1c, 1d). Using electrochromic shift (ECS) signals ^26^, we also followed the transition from State 2 to State 1 upon continuous illumination when PSII was inhibited (Fig. 1e). Thereby, we monitored the progressive reduction of PSI antenna size and, while this was seen in WT and T4A, a significant impairment was detected in T4E and ΔN (Fig. 1f). The ECS-based state transitions amplitudes in WT were comparable to those reported in literature ^27^, whereas T4A showed a reduced ability to revert to State 1 (Fig. 1g). The relative changes in both T4E and ΔN were the smallest.

**Fig. 1.**
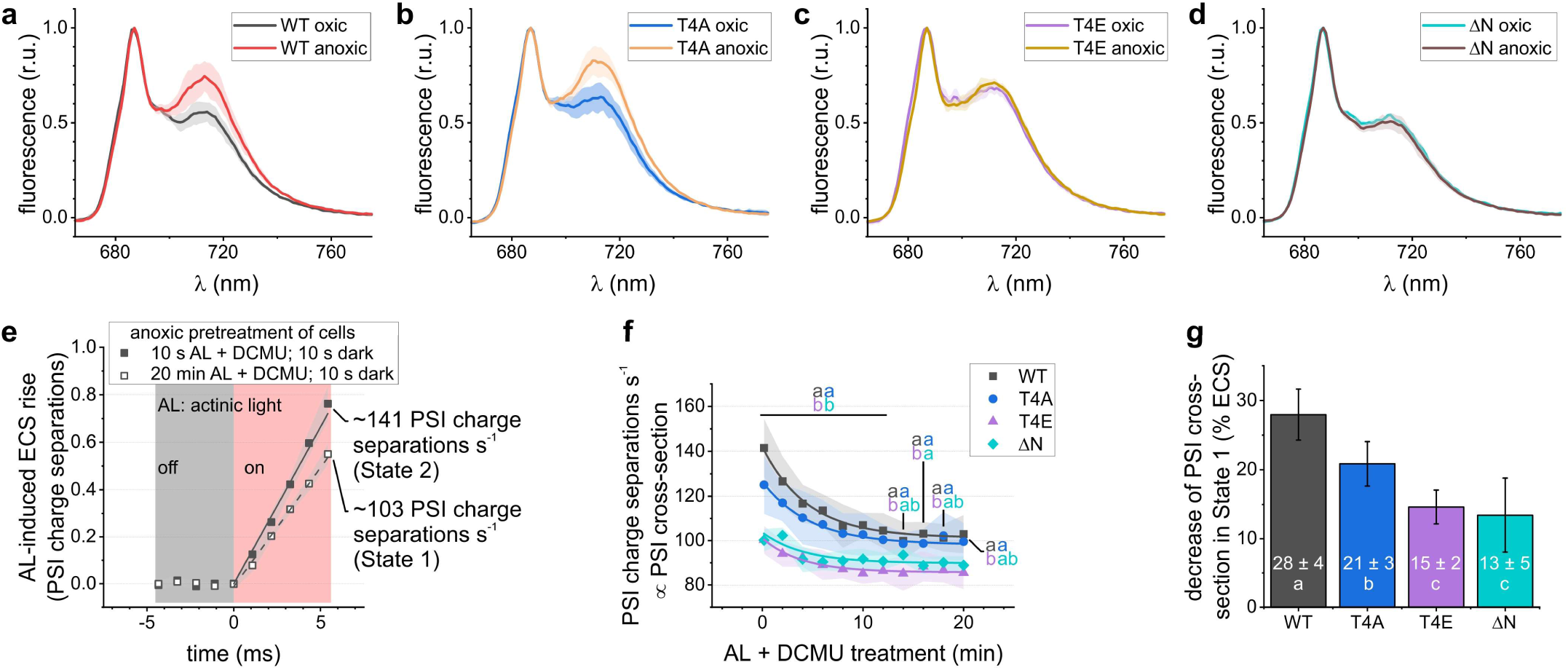
The relative antenna cross section is shown by measuring chlorophyll fluorescence emission spectra at 77 K and electrochromism. Panels **a** – **d** highlight the differences in antenna attached to photosystem (PS) I under State 1 (oxic) and State 2 (anoxic), contributing to ∼712 nm emissions. These spectra were averaged and normalized to PSII-attached antenna emission signals at ∼687 nm (biological replicates *N* = 3 ± SD). **e** The electrochromic shift (ECS) rise was recorded in DCMU-treated anaerobic cells during light-driven PQ pool oxidation. Traces were normalized to PSI. **f** The initial ECS slope decreased with reduction in PSI antenna size after state transition treatments. **g** The fitted amplitude from panel f was put in relation to the y-offset to calculate the extent of state transitions (*N* = 4 ± SD; letters indicate statistical significances via One-Way ANOVA/Fisher-LSD, *P* < 0.05).

The data in Fig. 1 suggest impaired STT7-dependent phosphorylation in both T4E and ΔN strains. To define specific STT7-dependent phosphorylation targets, we conducted phosphoproteomic analyses including a novel CRISPR/Cas9-generated *STT7* knockout mutant (Extended data Fig. 6). This *STT7* mutant served both as a benchmark reference for the PetD strains and as a tool to distinguish between effects stemming from different genetic *STT7* backgrounds (see also Extended data Table 1). To dissect STT7 function, cf. Fig. 2 and Extended data Fig. 2, as well as the following selection of results loosely categorized into five groups: (*i*) kinase autophosphorylation, (*ii*) LHCII and state transition complex formation, (*iii*) photoprotective CEF and NPQ, (*iv*) *b*_6_*f* and photosystem core subunits, and (*v*) miscellaneous phosphorylation targets.

**Fig. 2.**
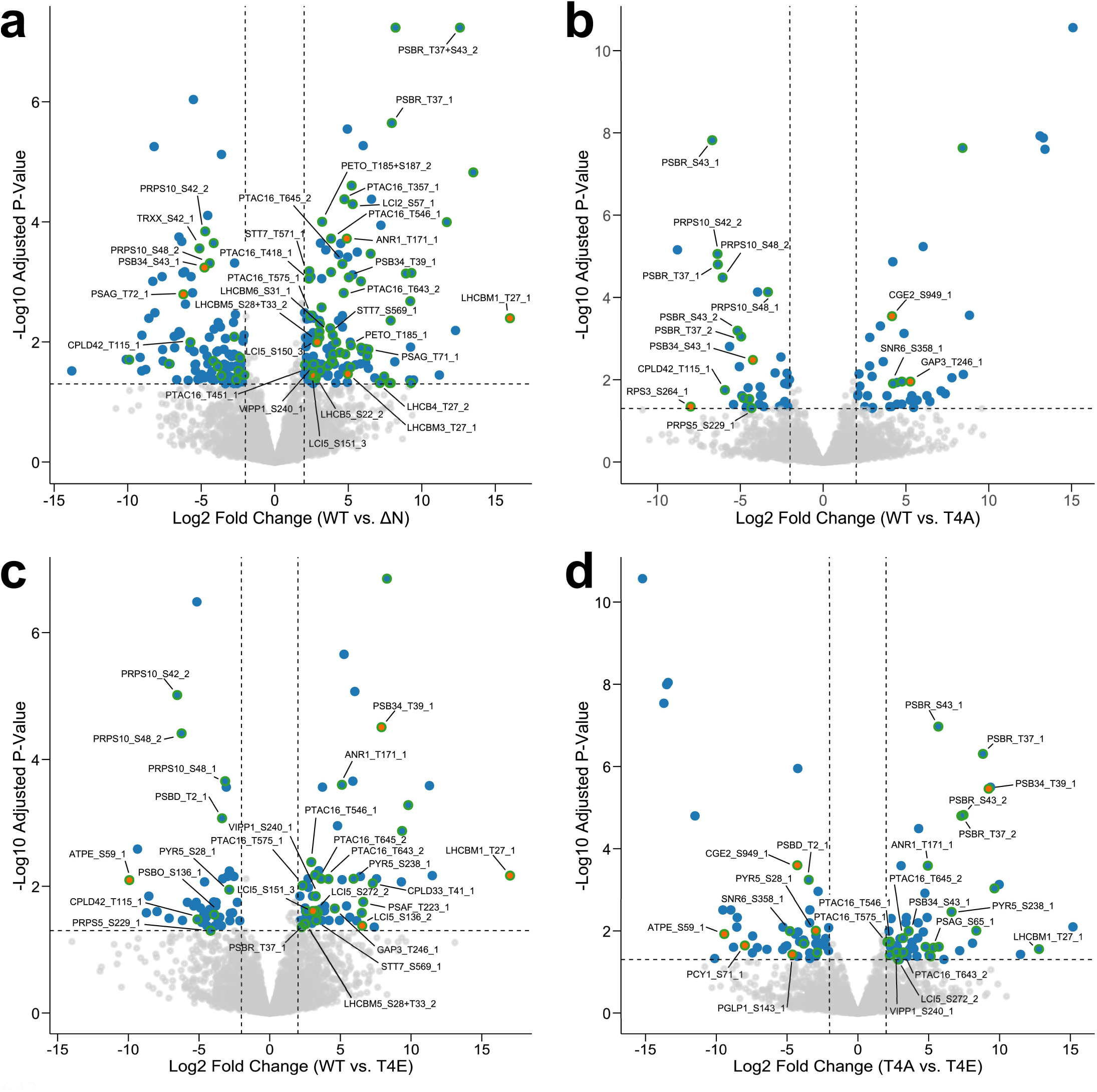
Volcano plots visualizing effect of PetD mutations on protein phosphorylation. Phosphorylation sites with FDR-adjusted p-value < 0.05 and |log_2_ fold change| > 2 are highlighted in blue, with threshold boundaries shown as dashed lines. Data points are shown in orange instead of blue when all values for one strain in the fold-change calculation are imputed due to completely missing data. Green halos indicate chloroplast-encoded proteins and nuclear-encoded proteins with confirmed or predicted (PB-Chlamy) chloroplast localization. Phosphorylation log_2_ fold changes are normalized to corresponding protein abundance levels to account for changes in protein expression. Numbers at the end of data labels indicate peptide multiplicity (total phosphorylations per peptide, e.g. PRPS10_S42_2 = PRPS10 with phosphorylated S42 and one additional phosphorylation on the same peptide). Comparisons are shown for **a** WT vs. ΔN, **b** WT vs. T4A, **c** WT vs. T4E, and **d** T4A vs. T4E.

An unexpected ^19^ observation was that, unlike *STT7*, the ΔN and T4E strains showed no detectable changes in LHCSR3 (T32) or LHCB4 double (T7+T11) phosphorylation. As these phosphorylation events are interrelated ^28^, this suggests either residual STT7 activity in ΔN and T4E, or a compensatory mechanism involving the physical presence of STT7 and possibly other chloroplast kinases such as STN8. Supporting the latter hypothesis, STN8 phosphorylation at T126 – absent in *STT7* – was retained in both ΔN and T4E, hinting at an intricate and potentially compensatory kinase network.

i. STT7 phosphorylation at S569/T571 was severely reduced in ΔN and T4E (Extended data Figs. 5b–5c), consistent with their failure to undergo state transitions. Nonetheless, STT7 autophosphorylation at T511, T513, T516, S533, and T547 remained largely unchanged across all strains (Extended data Figs. 5d–5h), suggesting that STT7 is partially functional in ΔN and T4E.
ii. In *STT7*, phosphorylation was broadly lost across key threonine clusters ^28^, notably the double phosphorylation in LHCB4 (T33+T35, T33+T37, T35+T37) and phosphorylation of T23 and T36 in LHCB5 (Extended data Table 1). LHCB5 retained only minimal phosphorylation at T20, and LHCB4 double-phosphorylation at T7+T11 was reduced by 6.8 log_2_-fold. Several background-independent ^19,29^ phosphorylation sites absent in *STT7* were also either absent (T27 of LHCBM1) or strongly diminished (double phosphorylation of S28+T33 in LHCBM5) in ΔN and T4E. The altered N-terminal phosphorylation in LHCBM1 and LHCBM5 likely explains the loss of state transitions in both strains, as these modifications are essential for assembly of the PSI–LHCI–LHCII state transition complex ^30^.
iii. As mentioned above, *STT7* lacked phosphorylation in the LHCSR3 N-terminal cluster (S26/S28/T32/T33/T39) ^28^, where up to three simultaneously phosphorylated residues were observed in WT. Meanwhile, LHCSR3 phosphorylation showed no significant difference between WT and the PetD variants. In contrast, ANR1 phosphorylation at T171 was confirmed to be a background-independent ^19,29^ target of STT7. It was also undetectable in ΔN and T4E. Notably, PETO levels were reduced nearly five-fold in ΔN (Extended data Fig. 3b). Given that PETO can chemically crosslink to K15 in the N-terminal region of PetD ^31^, its loss in ΔN and fg-loop mutants ^18^ supports a functional PetD–PETO interaction. Disruption of this interaction likely triggers PETO degradation.
iv. Consistent with prior data from an independent *STT7* mutant ^19^, PetD T4 phosphorylation was absent in *STT7*, while PSBD (T2) phosphorylation was markedly elevated (5.5 log_2_-fold). T4E exhibited a similar increase (3.4 log_2_-fold), whereas ΔN and T4A showed no significant changes at PSBD (T2). Both ΔN and T4E displayed reduced phosphorylation at PSB34 (T39), PSAG (T71), and PSBR (T37, S43, T37+S43), while ΔN specifically showed increased phosphorylation at PSAG T72. T4A, in contrast, exhibited a distinct profile: most phosphorylation sites were unchanged, but PSBR phosphorylation (T37, S43, T37+S43) was significantly elevated compared to the other strains.
v. PTAC16 – a component of the plastid transcriptionally active chromosome complex and a known STN7 substrate in *Arabidopsis* ^32,33^ – displayed the highest number of differentially phosphorylated sites. In *STT7*, PTAC16 phosphorylation was largely abolished. Similar patterns were observed in ΔN and T4E, though both retained some WT-specific phosphorylation. In contrast, the T4A mutant showed no significant changes in PTAC16 phosphorylation.

To assess functional consequences of the alterations in the engineered strains, we measured photosynthetic electron transfer rates (ETR) using ECS signals. The measured ETR in T4A and T4E were indistinguishable from WT, suggesting that phosphorylation at T4 does not have a direct impact (Figs. 3a, 3b). In contrast, ΔN cells showed a marked slowdown in ETR 30–50 ms after light onset under oxic conditions (Fig. 3a). Under anoxic conditions that favour CEF, ETR initially increased but reached a lower steady state, with the initial boost markedly reduced in ΔN compared to the other strains (Fig. 3b), indicating that the PetD N-terminus supports ETR via the *b_6_f*. Redox kinetics of P700 were also assessed (Figs. 3c, 3d). In ΔN, P700 oxidized rapidly upon illumination and re-reduced slowly in the dark, consistent with donor-side limitation under both oxic and anoxic conditions. Notably, steady-state ETR (Figs. 3a, 3b) were established despite substantial P700 acceptor-side limitation in all strains except in ΔN (Figs. 3c, 3d). This persistent donor-side limitation in ΔN was unaffected by the addition of the H⁺/K⁺ exchanger nigericin (Extended data Figs. 7a, 7b), excluding pH-dependent photosynthetic control. Treatment with DCMU and nigericin led to rapid P700 oxidation in all strains, slightly faster in ΔN (Extended data Figs. 7c, 7d), supporting impaired *b_6_f* function in both LEF and CEF.

**Fig. 3.**
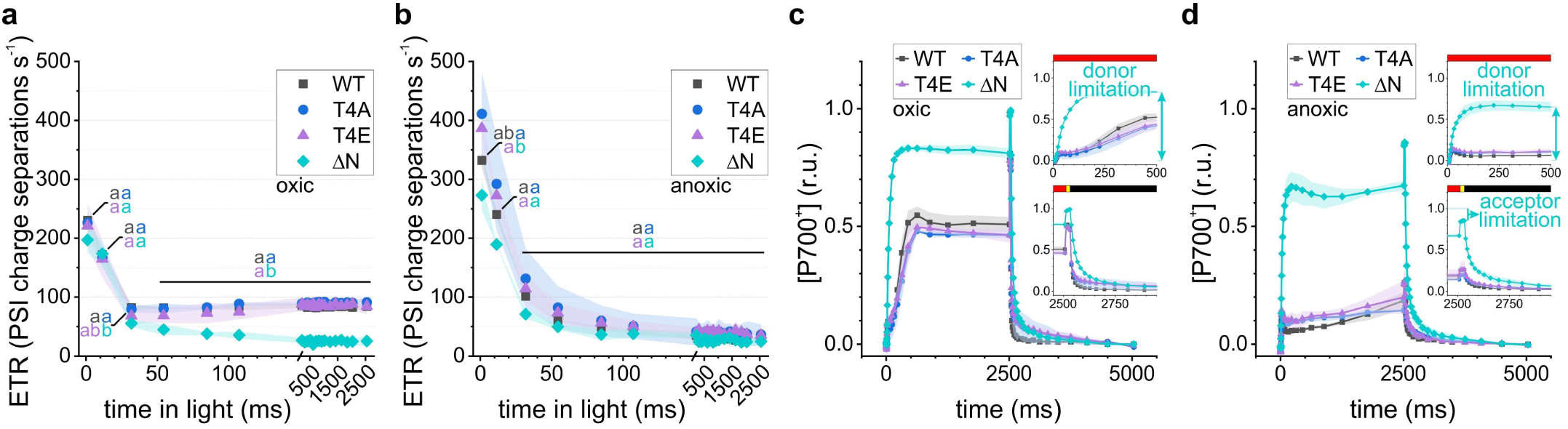
Photosynthetic activities of the cytochrome *b*_6_*f* variants are shown. Averaged kinetics are depicted (biological replicates *N* = 3 ± SD). **a**, Electron transfer rates (ETR) developed over the 2.5-s illumination period under oxic conditions, with the first detection at 1-ms of actinic light. The letters at the punctual rates indicate statistical significances (One-Way ANOVA/Tukey-HSD, *P* < 0.05) **b**, ETR developments are shown under anoxic conditions. **c**, Quantification of oxidized primary PSI donor ([P700^+^]) is plotted under oxic conditions. Red, yellow, and black bar represent actinic light intensities of 550, 3000, and 0 µmol photons m^-2^ s^-1^. **d**, The P700 kinetics are shown as in panel c but under anoxic conditions.

To confirm whether the observed differences ETR and P700 redox kinetics differences stemmed from *b_6_f* dysfunction, we evaluated *b_6_f* electrogenicity and redox kinetics (Extended data Fig. 8a, Fig. 4). ECS rise linked to Q-cycle activity was impaired exclusively in ΔN, despite the presence of heme *c*_i_ – the terminal electron acceptor in the low-potential chain ^34^ – as well as WT-like stability of the truncated *b_6_f* complex (Extended data Figs. 8b–c). Under oxic conditions (Fig. 4a), WT *b*-hemes showed a transient reduction (∼10 ms) before net oxidation (∼16 ms half-life), while cytochrome-*f* reduction (∼6 ms half-life) began ∼1 ms post-flash. In ΔN, the apparent *b*-heme reduction was enhanced because their oxidation slowed ∼20-fold (Fig. 4b). This coincided with a ∼10-fold slowdown in cytochrome-*f* reduction, indicating Qi-site impairment affecting the Qo-site as has been shown in *b_6_f* ^3,35^ and the related cytochrome-*bc*_1_ complex ^36,37^. Under anoxic conditions (Fig. 4c), WT displayed flat *b*-heme signals in the first 10 ms, followed by accelerated oxidation, consistent with prior findings ^3,35^. In ΔN, a redox-inactive low-potential chain caused a ∼25-fold slowdown in the high-potential chain, as reflected in cytochrome-*f* reduction (cf. Figs. 4c, 4d).

**Fig. 4.**
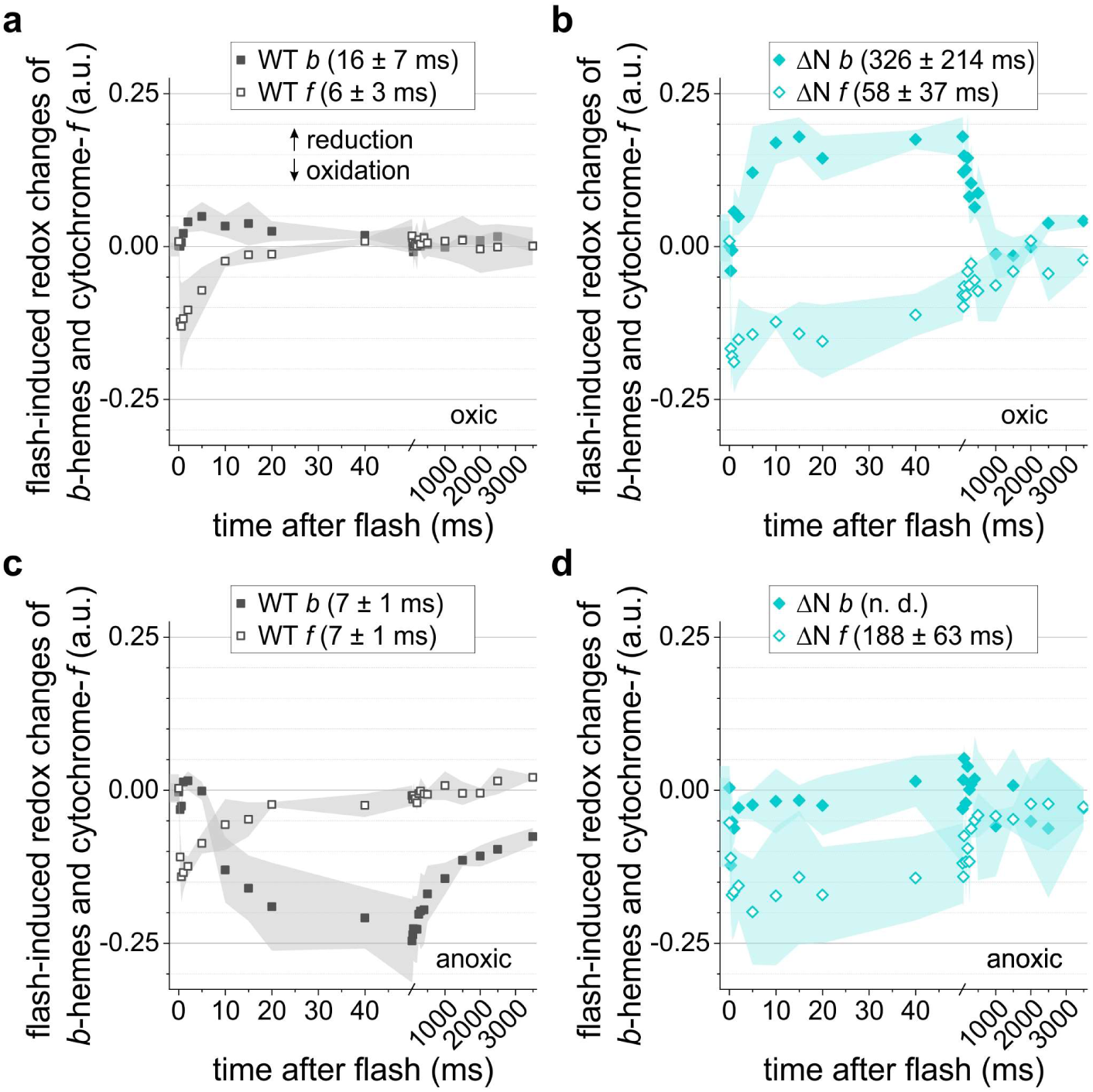
Single turnover redox changes of the cytochrome *b*_6_*f* hemes are shown. Averaged kinetics are depicted (biological replicates *N* = 3 ± SD). WT (**a**, **c**) as well as the ΔN mutant (**b**, **d**) were measured under oxic (**a**, **b**) and anoxic conditions (**c**, **d**). The averaged half-times of cytochrome-*f* reduction and *b*-hemes oxidation are given in the panels’ legends (n. d.; not determined).

The PetD N-terminus we investigated is exposed to the aqueous phase, contains two lysine residues, and is located near PetD D35. The latter is suggested to donate a proton to a water molecule in the Qi-site, which becomes displaced upon PQ binding ^38^. We propose that the ΔN truncation subtly affects the environment of the water channel or the functional properties of PetD D35. These changes may impair electron exchange in the low-potential chain, which could explain the pronounced slowdown in *b_6_f* turnover at both Qi- and Qo-sites (Fig. 4, Extended data Fig. 8a). This slowdown is consistent with the diminished ETR, enhanced P700 donor-side limitation (Fig. 3), and growth retardation observed in ΔN (Extended data Fig. 3a). Furthermore, diminished PQH_2_ oxidation at the Qo-site impedes STT7 activation ^39^, which may account for the partial inactivation of the kinase in the ΔN mutant (Fig. 2a). In contrast to ΔN, the phosphomimic T4E mutation does not affect *b_6_f* function. However, both mutations interfere with STT7 function, albeit through distinct and likely partial mechanisms: Qo-site defect in ΔN and negative charge introduction in the PetD stromal part at T4E.

To better understand the basis of STT7 inactivation, we analyzed its autophosphorylation pattern. Riche et al. ^18^ demonstrated via PhosTag PAGE that multiple STT7 autophosphorylation events are required for kinase activation, and that this process depends on binding of STT7 to *b_6_f*. Phosphorylation at residues S569 and T571 was strongly reduced in both ΔN and T4E strains, whereas other autophosphorylation sites, including T511/T513/T516, S533, and T547, were unaffected (Extended data Figs. 5d–5h). This data suggests that STT7 remains associated with the *b_6_f* complex in both mutant backgrounds but fails to undergo full activation. The partial autophosphorylation pattern lacking residues S569 and T571 likely contributes to the reduced kinase activity observed. However, we cannot exclude that STT7 binding to *b_6_f* is impaired in the two mutants.

Our findings point to a novel feedback mechanism in which STT7 activity is modulated by PetD T4 phosphorylation (Extended data Fig. 1). This feedback mechanism complements previously described STT7 autophosphorylation pathways involving other stroma-exposed structures of the *b_6_f* complex^17,18^. In some cases, such as PSBR, T4A mutants displayed elevated phosphorylation levels, consistent with disrupted feedback regulation. Two models may explain this phenomenon: (*i*) STT7 bound to *b_6_f* may detect PetD T4 phosphorylation and respond by reducing its own autophosphorylation, leading to decreased activity. (*ii*) Alternatively, STT7-dependent phosphorylation of PetD T4 could hinder subsequent STT7 binding to the *b_6_f* complex, thus indirectly limiting further kinase activation.

In conclusion, we show that the N-terminal region of PetD is critical for *b_6_f* function but not for complex stability, and that phosphorylation at PetD T4 regulates STT7 kinase activity via a novel autoregulatory feedback loop. This regulatory mode appears absent in organisms such as cyanobacteria^40^ and diatoms^41^, which lack PetD T4 and where state transitions are differently realized (Extended data Fig. 9). In Viridiplantae, feedback suppression of STT7 via PetD T4 may help coordinate state transitions with alternative regulatory mechanisms, including those targeting PSII core subunits such as PSBD T2 ^42,43^.

## Methods

### WT Plasmid construction

The *petD/trnR1* region (481 bp upstream and 1032 bp downstream of *petD*) was amplified from WT chloroplast DNA using restriction site-containing primer 1 (*SalI*) and 2 (*EcoRI*; for oligo sequences, see Extended data Table 1). The *SalI*/*EcoRI* digested PCR product was ligated into a double-digested pUC18 cloning vector and propagated as pUC18-3’-*petD*-5’ in NEB 5-alpha *E. coli* cells using ampicillin selection. In the next step, the aminoglycoside adenyl transferase (aadA) spectinomycin resistance cassette ^44^ was amplified from a pUC18-aadA vector using primers 3 and 4, each containing a *SpeI* restriction site. The cloned *petD/trnR1* region contains a single *SpeI* site into which the aadA cassette was cloned. To do so, intermediate vectors were created. First (pUC18-int1), the *SalI/SpeI* fragment from pUC18-3’-*petD*-5’ (1.46 kb) was ligated with the *SpeI* digested aadA PCR fragment, followed by PCR (primers 1 and 5) and blunt ligation of the ∼3 kb product into *SmaI* digested empty pUC18 vector. Second (pUC18-int2), the *EcoRI/SpeI* fragment from pUC18-3’-*petD*-5’ (0.51 kb) was ligated with the *SpeI* digested aadA PCR fragment, followed by PCR (primers 2 and 6), *XmaI*/*EcoRI* digest of the ∼2 kb product, and ligation into an empty pUC18 vector. The 2.4 kb fragment of *BamHI*/*BlpI* digested pUC18-int1 was ligated with the 3.7 kb fragment of *BamHI*/*BlpI* digested pUC18-int2 to yield pUC18-3’-*petD*-aadA-5’

### Point Mutation and Truncation Plasmid Construction

To generate T4E and T4A mutation as well as the *petD* truncation ΔN (lacking underlined N-terminal amino acids M<u>SVTKK</u>PD…), primers 8-11 and 13 were phosphorylated. Using pUC18-3’-*petD*-aadA-5’ as a template, five independent PCRs were done with the following primers: 1 and 8 (I), 1 and 9 (II), 1 and 10 (III), 11 and 12 (IV) and 12 and 13 (V). Ligated PCR fragments I and IV served as templates for the T4E PCR (primers 1 and 12) which was further digested (*PacI*/*SalI*) for ∼1.36 kb insert swapping and mutant introduction into pUC18-3’-*petD*-aadA-5’ (ligation with ∼4.78 kb backbone). In a similar fashion, for T4A PCR (primers 1 and 12) PCR fragments II and IV served as templates. For ΔN PCR (primers 1 and 12), fragments III and V served as templates.

### Biolistic Transformation of WT Strain

Mutant strains were generated through chloroplast transformation using plasmids containing the modified *petD* gene version along with an *aadA* cassette to confer spectinomycin resistance (150 µg/ml). The recipient wild-type (WT) strain carries a six-histidine tag at the C-terminus of PetA ^21^. Spectinomycin resistant clones were selected and replated several times to obtain homoplastic strains (for sequencing primers, see Extended data Table 1).

### Growth conditions and treatments

Cells were maintained at 50 µmol photons m^-2^ s^-1^ at 23°C on agar plates of Tris-acetate-phosphate (TAP) medium or Tris-minimal medium (TP) in the absence of acetate ^45^ by bubbling sterile air under a 16-h light/8-h dark regime. For state transition treatment using 77K fluorescence, TAP-grown synchronized cultures were freshly diluted one day before the experiment and, upon harvesting (4000 *g*, 5 min, 23°C), adjusted to 5 µg chlorophyll ml^-1^. Aliquots were either kept for 45 min in aerobic conditions by shaking in 40 µmol photons m^−2^ s^−1^ light upon adding 10 µM 3-(3,4-Dichlorophenyl)-1,1-dimethylurea (DCMU; State 1), or were given anoxic treatment in the dark using a glucose oxidase/catalase cocktail (State 2) ^46^. For proteomic analyses of *STT7* experiments, TAP grown cells were shifted to State 2 conditions for 30 min and samples for phospho-proteomic analyses were collected. For ECS-based state transition measurements, the cells (40 µg chlorophyll ml^-1^) were bubbled with argon gas for 15 min in the dark, followed by 1:1 dilution in TP-Ficol (20% *w*/*v*) and addition of 20 µM DCMU. The strains were checked for photoautotrophic growth for two weeks by spotting 20 µl of 3×10^5^ cells/ml (and dilutions) on TP agar plates at 25°C kept under continuous light at 40 and 200 µmol photons m^−2^ s^−1^.

### Biochemical methods

Liquid cultures of WT, T4A, T4E, and ΔN were grown in TAP medium and freshly diluted one day before harvesting. The cells were pelleted and mixed in a buffer containing 0.2 M dichloro-diphenyl-trichloroethane, 0.2 M Na_2_CO_3_, 10 mM NaF, and protease inhibitors (0.2 mM phenylmethylsulfonyl fluoride, 1 mM benzamidine, 20 mM amino caproic acid) to be analyzed via SDS-PAGE ^47^. For heme staining the *b_6_f* was isolated using the standard protocol ^48^ to analyze their purity via SDS-PAGE. The high-spin hemes were stained in gel using 3,3’,5,5’-tetramethylbenzidine ^49^. Blue Native (BN) PAGE was carried out according to the manufacturer’s protocol (NativePAGE Bis-Tris; Thermo Fisher Scientific)

### Generation of *STT7* mutants with CRISPR-CAS9

Chlamydomonas strain CC-125 (*nit1*; *nit2*; *mt+*) was grown in tris-acetate-phosphate (TAP) media under continuous illumination of 50 μmol photons m^−2^ s^−1^ at 22 °C. Single guide RNAs (sgRNAs) were designed by CHOPCHOP using version 5.6 of the *Chlamydomonas reinhardtii* genome, and 5’-CTCCTGTAGACCGCTCAATGCGG was selected as the targeting sequence, and sgRNAs were bought will all accessory parts to make it functional (IDT, USA). Ribonucleic protein (RNP) complexes were prepared by duplexing the sgRNA (23% volume per volume, v/v; IDT) and Cas-9 (19% v/v; IDT) within a duplex buffer (58% v/v; IDT) and incubated at 37°C for 25 to 30 min. After treatment with autolysin for 3 hours without shaking, cells were washed with TAP media supplemented with 40 mM of sucrose, resuspended with the same supplemented media to approximately 2 x 10^8^ cells mL^-1^, and electroporated in the presence of a hygromycin resistance DNA cassette (2 µg mL^-1^) and the 37°C-incubated RNP mixture. After 10 min at room temperature, cells were resuspended in 10 mL of TAP media supplemented with 40 mM of sucrose and left without shaking overnight in dim light (10-20 µmol photons m^-2^ s^-1^) at 22°C. Hygromycin resistant transformants were selected on solid TAP media (2% Agar w/v) containing hygromycin (20 µg mL^-1^). sgRNA targeted regions were amplified via polymerase chain reaction (PCR) to check for full or partial insertion of the hygromycin resistance cassette at the target site using the following primers: STT7 F/R (for oligo sequences, see Extended data Table 1) with an annealing temperature of 62.6°C (Extended Data Fig. 6). Impaired accumulation of the STT7 protein was then verified by immunodetection (Agrisera, AS15 3080).

### Sample preparation for mass spectrometry

Protein isolation from cell pellets was carried out as described ^43^. Reduction/alkylation of cysteines and tryptic digestion of proteins (100 µg per sample) was performed following the SP4 protocol ^50^. After digestion, peptides were desalted using PurePep H50 SPE Spin columns (Affinisep). The peptide samples were divided into two aliquots: 5 µg for whole proteome analysis and 95 µg for phosphopeptide enrichment. Both aliquots were dried using vacuum centrifugation before further processing.

To enrich phosphopeptides, titanium dioxide (TiO2) beads with 5 µm particle size (Titansphere, GL Sciences) in a metal oxide affinity chromatography (MOAC) procedure were used. TiO2 (1 mg per sample) was activated once with acetonitrile and equilibrated three times with loading buffer (LB; 80% (v/v) acetonitrile/5% (v/v) TFA/1 M glycolic acid). Loading buffer was added to create a 10% (w/v) TiO2 suspension. Peptide samples were dissolved in 50 µl LB, combined with 10 µl TiO2 suspension, and incubated for 30 minutes at 25°C in an Eppendorf Thermomixer at 1,200 rpm). The mixture was transferred to SDB-XC STAGE tips prepared in-house ^51^. Unless stated otherwise, all subsequent washing and elution steps were performed using 60 µl buffer volumes with in-between centrifugation at 2000 × g at room temperature. The buffer was removed by centrifugation and the TiO2 beads were washed twice with LB and once with 1% (v/v) acetonitrile/0.1% (v/v) TFA (W1). Non-phosphorylated peptides that were removed from the beads and subsequently bound to the SDB-XC membrane plug were removed by one wash with 80% (v/v) acetonitrile/0.1% (v/v) TFA (W2). The phosphopeptides were eluted from the TiO2 beads onto the SDB-XC material using 160 µl of pH 11 buffer (0.4 M sodium hydrogen phosphate/NaOH with 1% acetonitrile). After washing once with 1% acetonitrile/0.1% TFA, the phosphopeptides were sequentially eluted in three fractions using increasing concentrations of acetonitrile (7.5%, 20%, and 60%) in 100 mM ammonium formate (pH10). For samples from the STT7 experiment, only one fraction (60% acetonitrile in 100 mM ammonium formate (pH10)) was collected. The strongly bound phosphopeptides were eluted in two steps: first from the TiO2 beads using 5% ammonia, then from the SDB-CX membrane using 60% acetonitrile in 100 mM ammonium formate at pH 10. All fractions were dried by vacuum centrifugation and stored at −80°C until MS analysis.

### Mass spectrometry

#### Whole proteome analysis

Dried peptide samples were reconstituted in 5 µl of 2% (v/v) acetonitrile/0.05% (v/v) TFA in LC/MS grade water. Samples were analyzed on an LC-MS/MS system consisting of an Ultimate 3000 NanoLC (Thermo Fisher Scientific) coupled via a Nanospray Flex ion source (Thermo Fisher Scientific) to a Q Exactive Plus mass spectrometer (Thermo Fisher Scientific). Peptides were concentrated on a trap column (Acclaim Pepmap C18, 5 × 0.3 mm, 3 µm particle size, Thermo Scientific) for 3 min using 4% (v/v) acetonitrile/0.05% (v/v) in LC/MS grade water at a flow rate of 10 µl/min. The trap column was operated in back-flush mode, allowing the transfer of peptides on a reversed-phase column (PepSep Fifty, 500 × 0.075 mm, 1.9 µm particle size, Bruker) for chromatographic separation. The eluents used were 0.1% (v/v) formic acid in LC/MS grade water (A) and 0.1% (v/v) formic acid/80% (v/v) acetonitrile in LC/MS grade water (B). The gradient was programmed as follows: 2.5% to 5% B over 5 min, 5% to 17.5% B over 47 min, 17.5% to 40% B over 105 min, 40% to 99% B over 10 min, and 99% B for 20 min. Flow rate was 250 nl/min. The mass spectrometer was operated in data-dependent acquisition (DDA) mod, alternating between one MS1 full scan and up to 12 MS2 scans. Full scans were acquired with the following settings: AGC target 3e6, MS1 resolution 70000, maximum injection time 50 ms, scan range: 350–1400 m/z. Settings for MS were: AGC target 5e4, resolution 17500, MS2 maximum injection time 80 ms, intensity threshold 1e4, normalized collision energy 27 (HCD)). Dynamic exclusion was set to ‘auto’, assuming a chromatographic peak width (FWHM) of 60 s.

### Phosphoproteomic analysis

Prior to analysis, dried peptide samples were reconstituted in 5 µl of 2% (v/v) acetonitrile/0.05% (v/v) TFA in LC/MS grade water. LC-MS/MS system, flow rate and eluent compositions were the same as described above. The gradient for peptide separation was as follows: 2.5% to 5% B over 5 min, 5% to 40% B over 92 min, 40% to 99% B over 10 min, and 99% B for 20 min.

The mass spectrometer used the same DDA mode settings as the whole proteome analysis, with two changes: MS2 maximum injection time was increased to 120 ms, and dynamic exclusion was set to 45 s.

### MS data analysis

For quantitative phosphoproteome analysis of four biological replicates, mass spectrometry raw data from nonenriched and enriched samples were searched using Fragpipe ^52,53^ against a concatenated, non-redundant database containing *Chlamydomonas reinhardtii* protein sequences from v5.6 and v6.1 gene models (Phytozome 13, phytozome-next.jgi.doe.gov). In addition, the polypeptide sequences of mutated/tagged proteins were added to the database. Default ‘LFQ-MBR’ workflow settings were applied to data from whole proteome samples, while the ‘LFQ-phospho’ workflow was used for phosphopeptide samples. A false discovery rate of 1% was applied at both peptide and protein levels.

Fragpipe output files containing peptide/protein identifications and quantitative data (’combined_modified_peptide.tsv’ from enriched samples and ‘combined_protein.tsv’ from whole proteome samples) were reformatted using a custom Python script to ensure compatibility with Phospho-Analyst (v. 1.0.2), which was used for differential expression analysis ^54^: Missing values were imputed using the ‘MinProb’ function, and variance stabilizing transformation was applied to normalize phosphosite intensities. Since the whole proteome LFQ data was already normalized in Fragpipe, no additional normalization steps were needed. Phospho-Analyst automatically corrected phosphosite abundance changes for underlying protein levels. A phosphorylation site localization probability threshold of 0.33 was applied, and p-values were adjusted using the Benjamini-Hochberg method. Predicted protein localizations are based on recently published data (PB-Chlamy)^55^.

### Optical spectroscopy

The flash-induced *b_6_f* measurements were carried out by using a Joliot-type spectrophotometer (JTS-150, SpectroLogiX) equipped with the interference filters and deconvolution procedures as reported previously ^3^. Cells were adapted to alternating 7.5 s darkness and 5 s actinic light (AL; 550 µmol photons m^-2^ s^-1^ peaking at 630 nm). Sub-saturating 6-ns flashes (Q-switched Nd:YAG, Continuum) were fired at 3 s darkness, hitting ∼40% of PSI. Three averaged baseline spectra (in the absence of a flash) were subtracted from each flash measurement at the *b_6_f*-specific wavelengths. For multiple turnover conditions, previously described conditions and protocols used cells which were adapted to alternating 2.5 s AL and dark. Briefly, ETR were obtained in the presence of LEF but referred to PSI charge separation activity. This stems from separate ECS calibration measurements that allowed, via saturating single-turnover flashes, for the conversion of optical changes (obtained as Δ*I*/*I* at 520 nm - 546nm) into the ECS signals corresponding to 1 PSI charge separation ^56^. The ETR protocol was based on a Dark Interval Relaxation Kinetics approach ^57,58^. P700 measurements relied on the Klughammer & Schreiber method ^59^. The ECS-based state transition method is described elsewhere ^26^; it measures PSI-driven charge separation rates as a function of PSI antenna size. Accordingly, the extent of State 2 to State 1 transition under light and DCMU displayed as a flattening of ECS slopes acquired during a 20-min measurement sequence (AL as above, 10 µM DCMU; Figures 1e–g). The kinetics were fitted using the exponential decay function 𝑦 = 𝑦_0_ + 𝐴𝑒*^−(x-x_0_)∕tl^*. The state transition extent was expressed as (𝐴 + 𝑦_0_)/𝑦_0_, where 𝑦_0_ is the y-offset and 𝐴 the amplitude. For fitting, the x-offset *x*_0_ was fixed at 10 s, corresponding to the minimal pre-illumination period of the State 2 reference sample.

## Data availability statement

All data supporting the findings of this study are available within the paper and its Supplementary Information. The mass spectrometry proteomics data have been deposited to the ProteomeXchange Consortium (proteomecentral.proteomexchange.org) via the PRIDE partner repository ^60^ with the dataset identifier PXD060640. The generated mutant cell lines are available on request from authors. The *STT7* mutant is available in the Chlamydomonas Resource Center (CC-6272; www.chlamycollection.org).

## Acknowledgements

M.H. (507704013) and F. B. (507704013) acknowledge funding by the Deutsche Forschungsgemeinschaft (DFG FOR 5573/1). M.H. acknowledges the RECTOR program in association with the University of Okayama, Japan. F. B. acknowledges Deutsche Forschungsgemeinschaft 461765884. Y.M. acknowledges the Alexander von Humboldt Foundation for a Research Fellowship for postdocs (1219125). A.B. acknowledges the support of Carnegie Science. The authors thank Sai Kiran Madireddi for helping with the initial production and characterisation of the *STT7* mutant.

## Author contributions

M.H. and F.B. designed the project. A.Z. and F.B. did genetic engineering and kinetic measurements and analyses. S.B. contributed to mutant analyses. C.S. and A.B. produced the *STT7* KO strains. M.S. did the mass spectrometric measurements and data evaluation. M.S. and M.H. analysed the mass spectrometric data. M.H wrote the manuscript with contributions from F.B., A.Z., A.B. and M.S. All authors contributed to the analysis and the final version of the manuscript.

---

**Extended Data Fig. 1.**
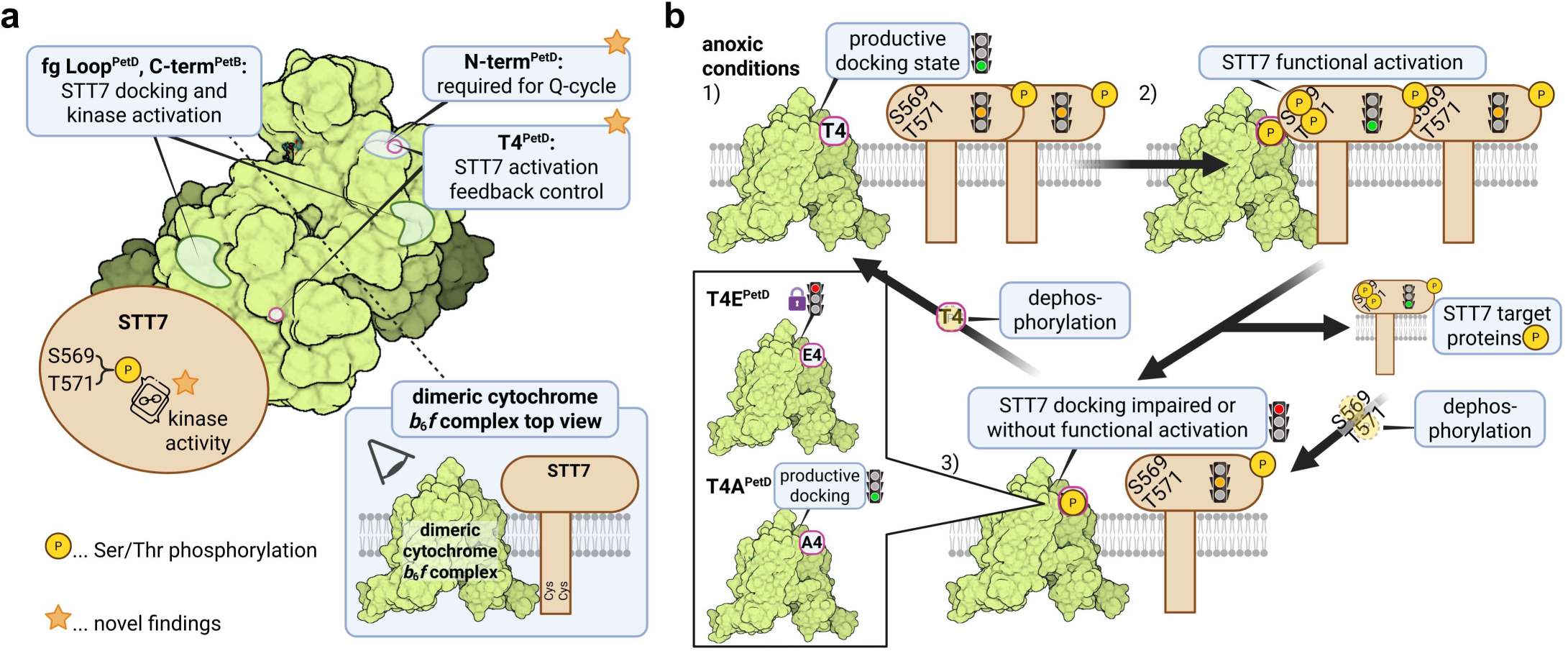
Graphical abstract of the cytochrome *b*_6_*f* complex interplay with STT7 kinase regulation. **a**, The N-terminus of PetD has been investigated and the focus was on T4. The model is built on previous STT7 docking studies (references 17 and 18 in the main text) and disregards possible STT7 oligomerization and redox regulation for the sake of simplicity. **b**, The feedback loop upon PetD T4 phosphorylation as well as investigated mutants is summarized. The figure was created with Biorender.

**Extended Data Fig. 2.**
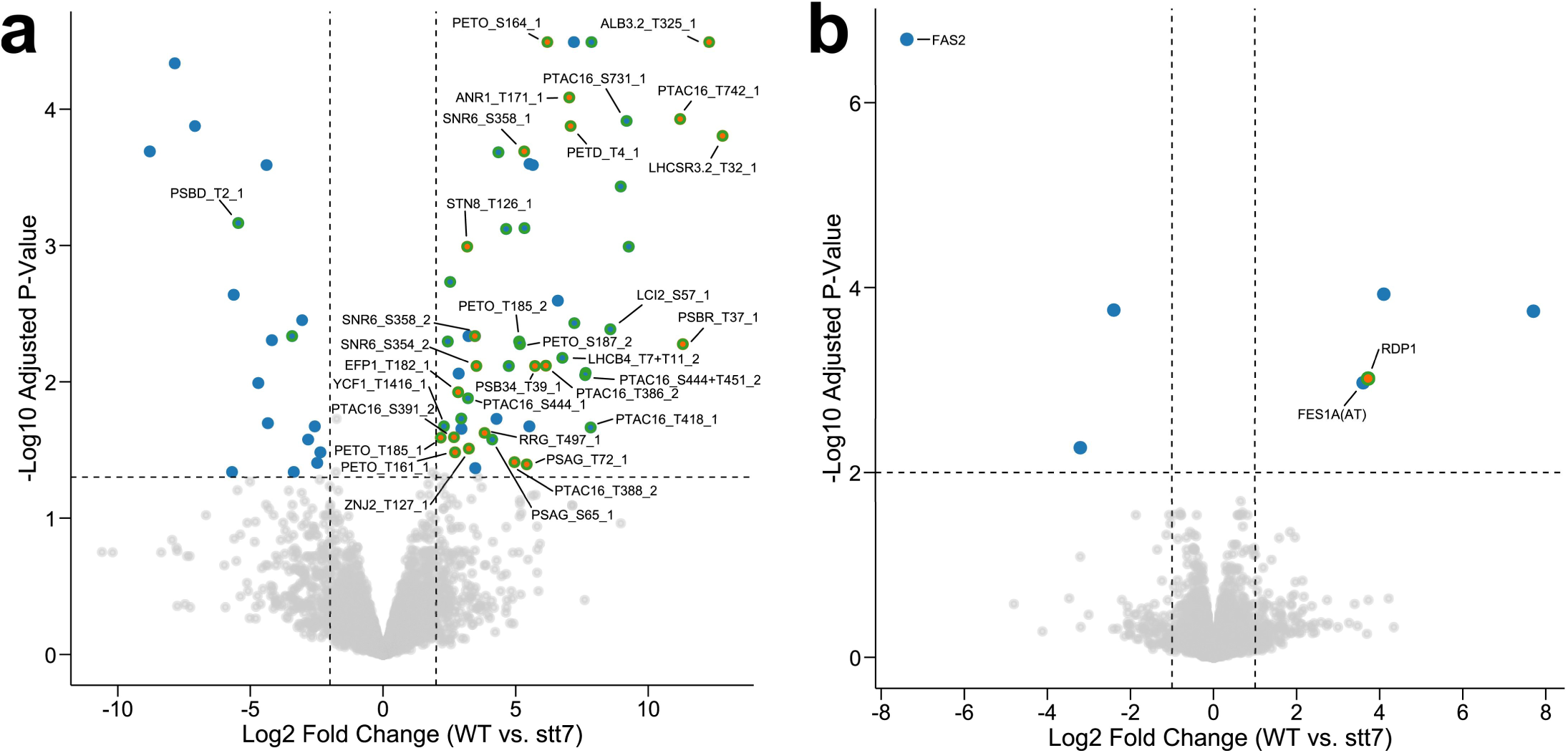
Volcano plots visualizing differential phosphorylation (a) and protein expression (b) between WT and *STT7* mutant. Proteins with FDR-adjusted p-value < 0.01 and |log_2_ fold change| > 1 are highlighted in blue, with threshold boundaries shown as dashed lines. Data points are shown in orange instead of blue when all values for one strain in the fold-change calculation are imputed due to completely missing data. Green halos indicate chloroplast-encoded proteins and nuclear-encoded proteins with confirmed or predicted (PB-Chlamy) chloroplast localization.

**Extended Data Fig. 3.**
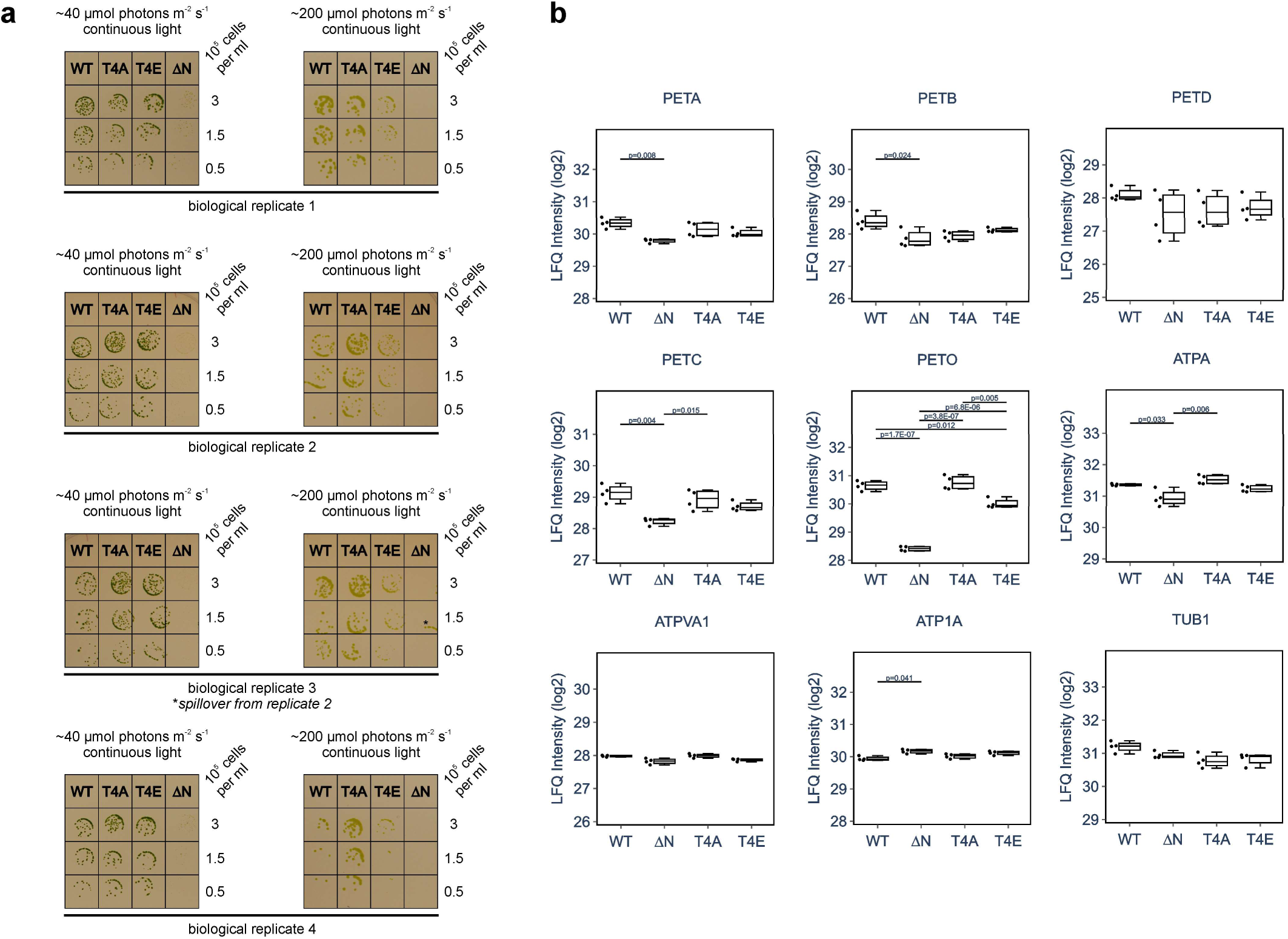
Growth analysis in varying light conditions and protein expression in the mutants. **a**. Spot tests on minimal media compare the growth under two light conditions (40 and 200 µmol photons m^-2^ s^-1^). **b**. Protein expression of *b_6_f* subunits and control proteins in the engineered PetD strains. The box plots visualize the log₂ protein LFQ intensities.

**Extended Data Fig. 4.**
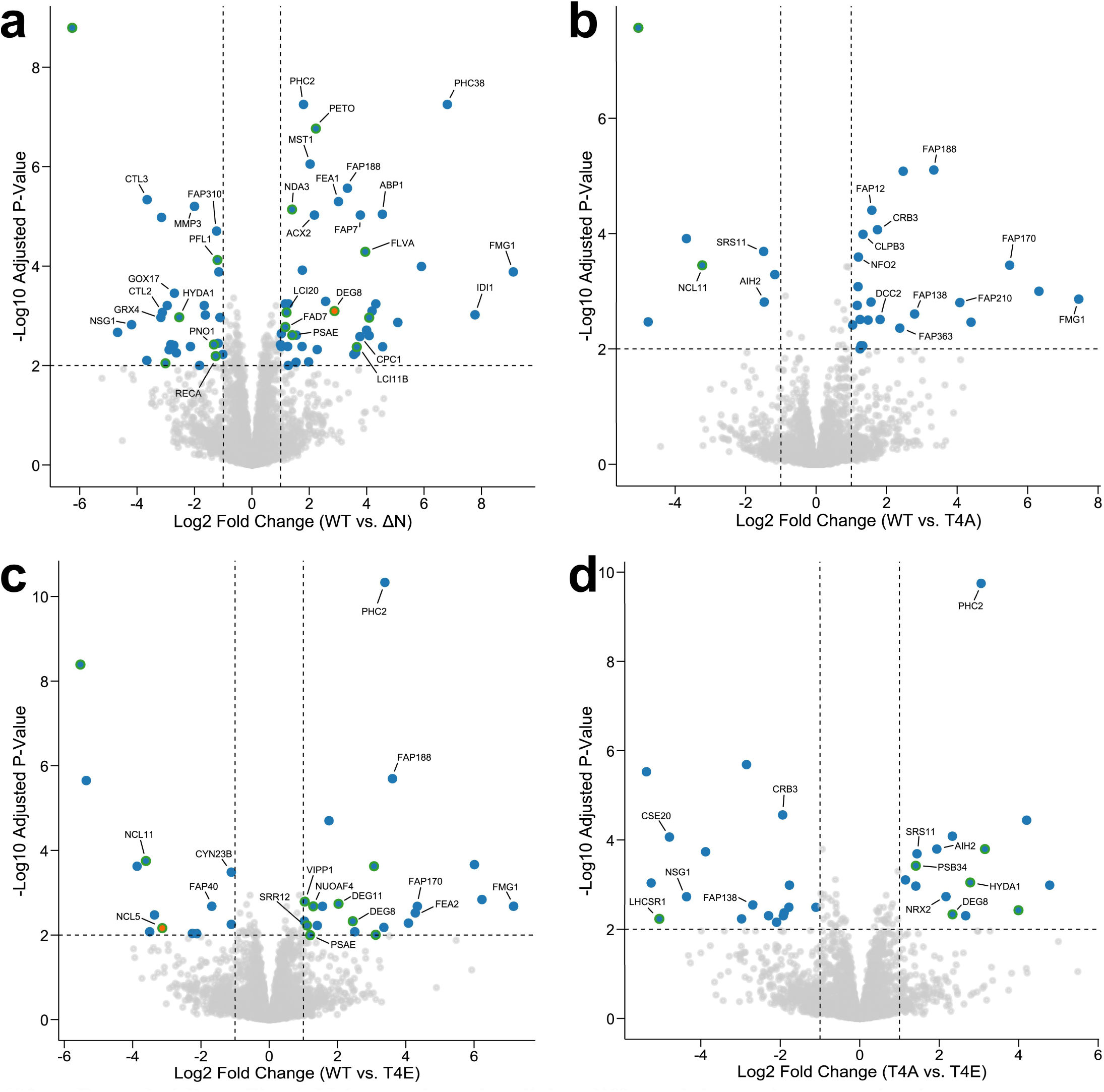
Volcano plots visualizing differential protein expression between WT and PetD mutants. Proteins from four biological replicates with FDR-adjusted p-value < 0.01 and |log₂ fold change| > 1 are highlighted in blue, with threshold boundaries shown as dashed lines. Data points are shown in orange instead of blue when all values for one strain in the fold-change calculation are imputed due to completely missing data. Green halos indicate chloroplast-encoded proteins and nuclear-encoded proteins with confirmed or predicted (PB-Chlamy) chloroplast localization. Comparisons are shown for **a** WT vs. ΔN, **b** WT vs. T4A, **c** WT vs. T4E, and **d** T4A vs. T4E.

**Extended Data Fig. 5.**
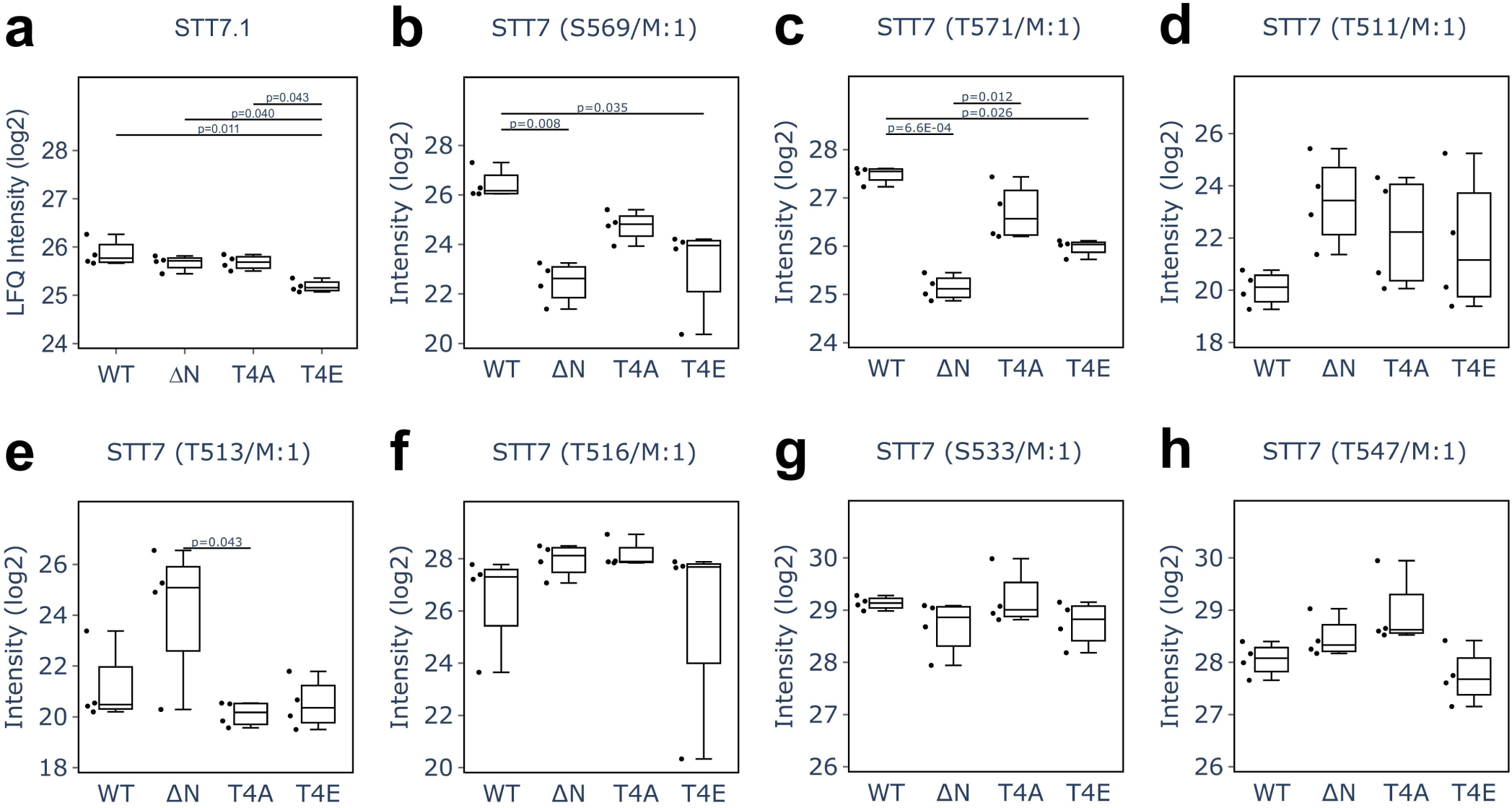
STT7 protein expression and selected autophosphorylation amounts in the engineered PetD strains. The box plots visualize significant differences of log_2_ fold changes in (**a**) protein LFQ and (**b**, **c**) phosphosite intensities from four biological replicates. No significant intensity differences were detectable for various STT7 phospho-peptides (panels **d** – **h**).

**Extended Data Fig. 6.**
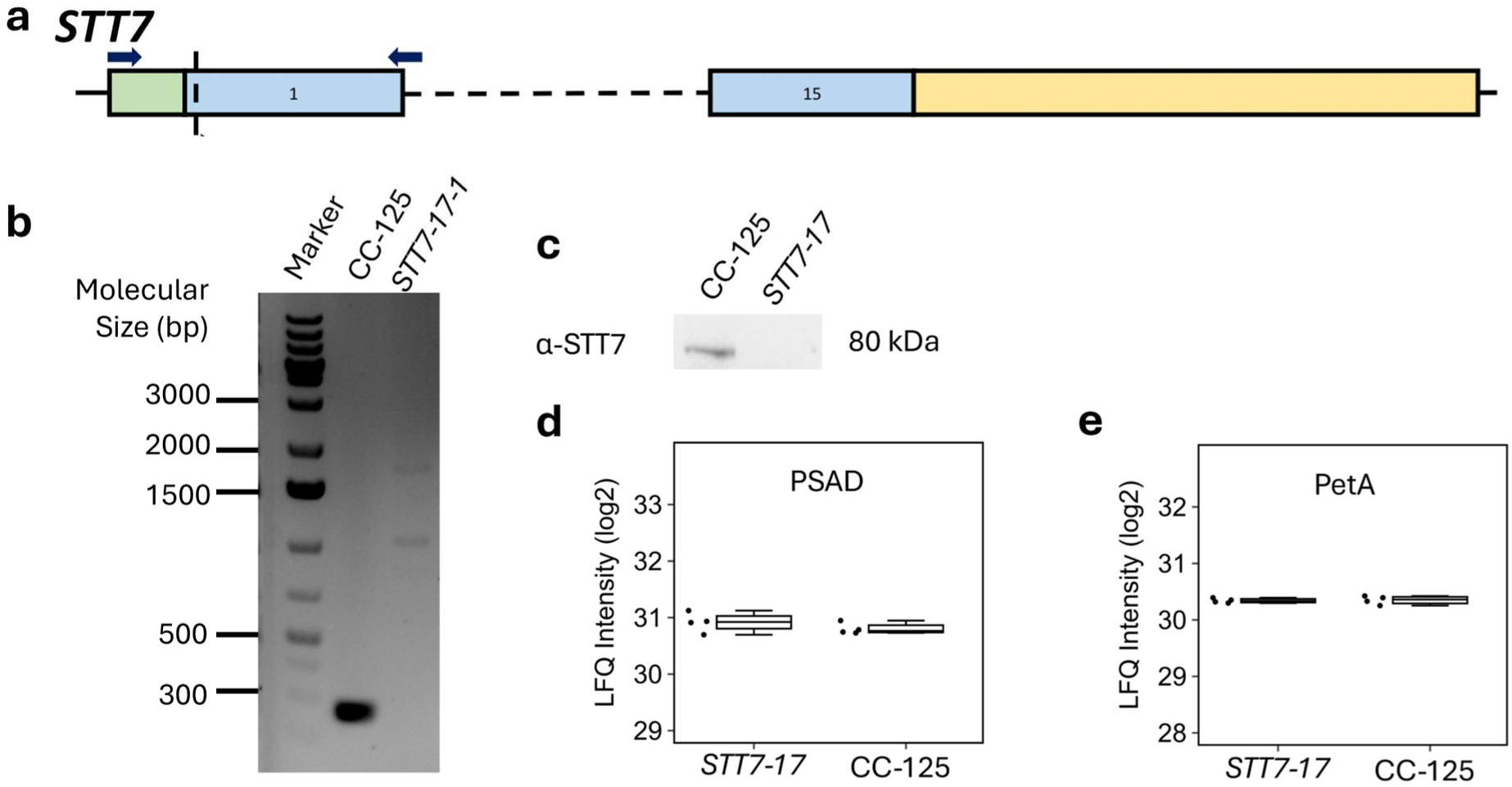
Generation of the *STT7* CRISPR/Cas mutant. **a.** The gene map of the *STT7* gene showing 5’ UTR (green), numbered exons (blue) and 3’UTR (yellow) with predicted cut site (vertical dotted line) and primers used to verify the genotype of the insertion (blue arrows). **b.** Genetic characterization of the *STT7* mutant and its control (CC125) using PCR on genomic DNA. **c.** Immunodetection of STT7 in the *STT7* mutant and its control (CC125). The box plots in panels **d** and **e** visualize the log₂ protein LFQ intensities of PSAD and PetA, respectively.

**Extended Data Fig. 7.**
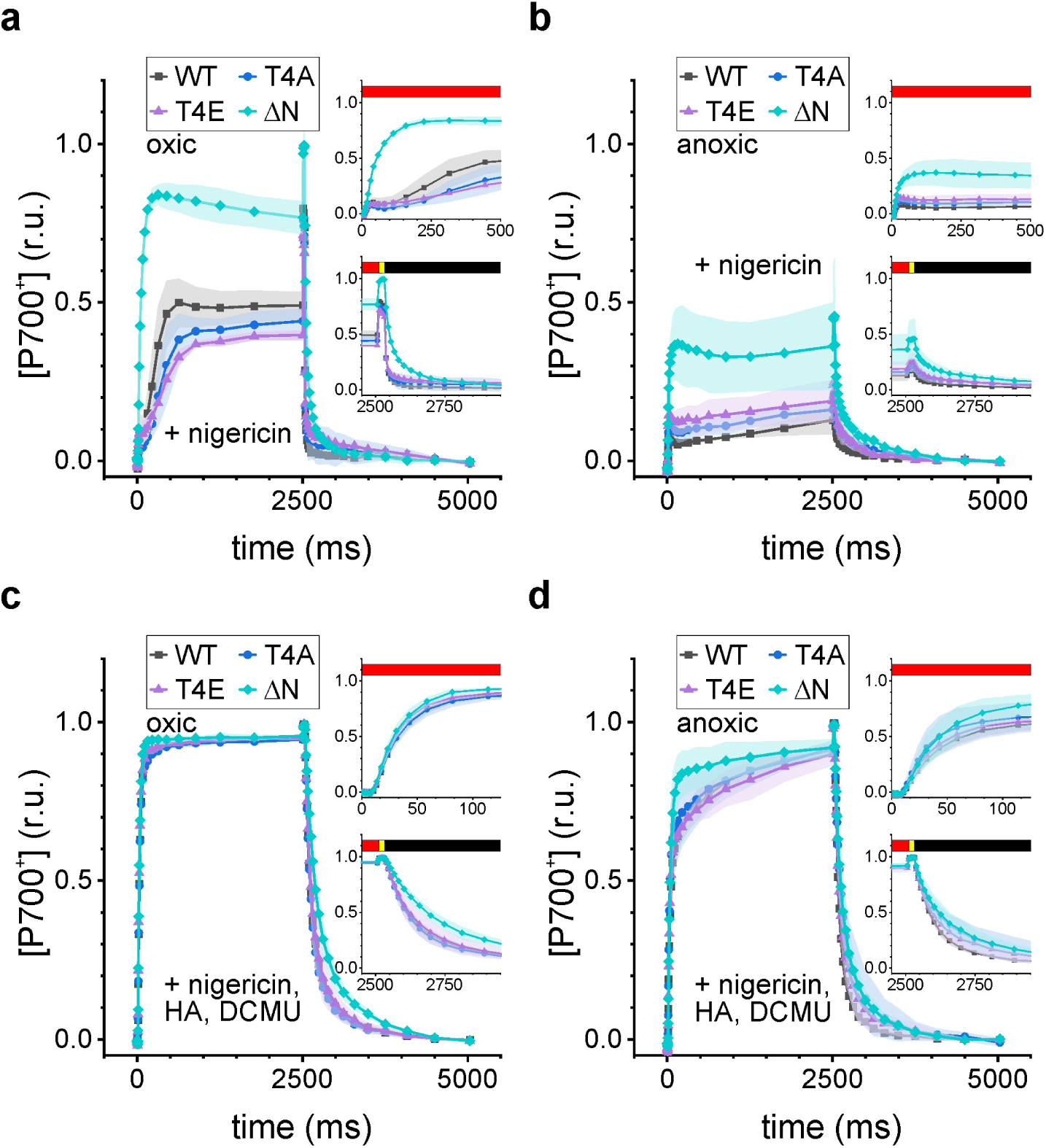
Photosynthetic activities of the cytochrome *b*_6_*f* variants are shown for P700. Averaged kinetics are depicted (biological replicates *N* = 3 ± SD) and the reference measurements are shown in main text Figs. 3c and 3d. The redox changes of primary PSI donor ([P700^+^]) is plotted (**a**) under oxic and (**b**) anoxic conditions in the presence of 10 µM nigericin, acting as H^+^/K^+^ exchanger to attenuate ΔpH in favor of an electric field. Red, yellow, and black bar represent actinic light intensities of 550, 3000, and 0 µmol photons m^-2^ s^-1^. **c**, The conditions were as in panel a but photosystem II was inhibited by hydroxyl amine (HA; 2 mM) and 3-(3,4-Dichlorophenyl)-1,1-dimethylurea (DCMU; 20 µM). **d**, The conditions were as in panel c except for the anoxic pre-treatment.

**Extended Data Fig. 8.**
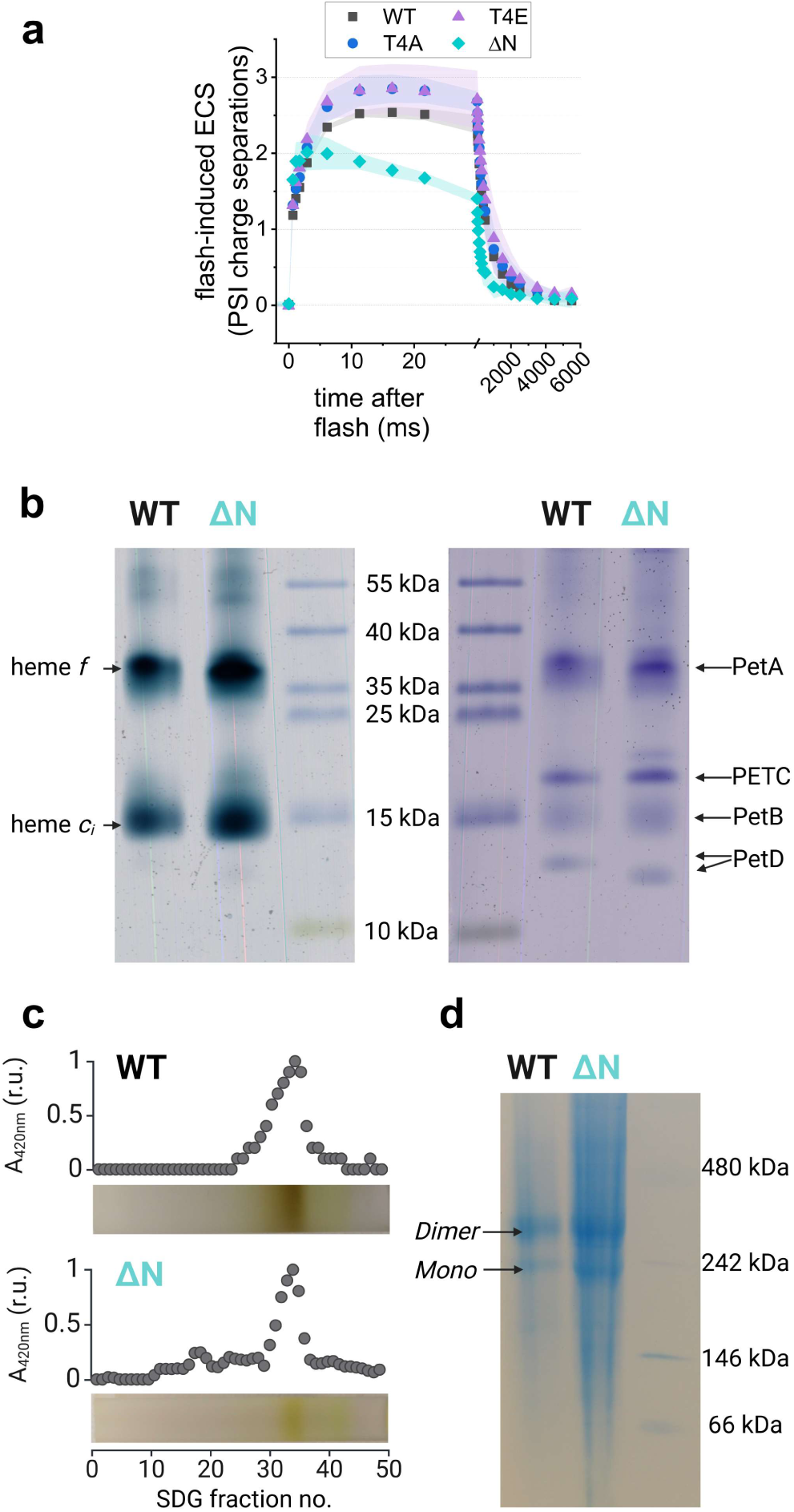
Laser flash-induced electrochromic shift (ECS) measurements show striking differences in ΔN and are not related to stability properties mutant *b*_6_*f*. **a.** The ECS shows multiphasic kinetics that rely on activities of both photosystems, the cytochrome *b*_6_*f* complex and ATP synthase. Averaged measurements are shown (biological replicates *N*=3 ± SD). **b.** Heme *c*_i_ was present in the ΔN mutant strain, as shown by in-gel heme staining with TMBZ. Coomassie-stained gels of 10 µg *b*_6_*f* are shown for reference. **c.** Sucrose density gradient (SDG) fractionation and **d.** BN-PAGE analysis of 5 µg pooled *b*_6_*f* showed no increased monomerization tendency in ΔN particles.

**Extended Data Fig. 9.**
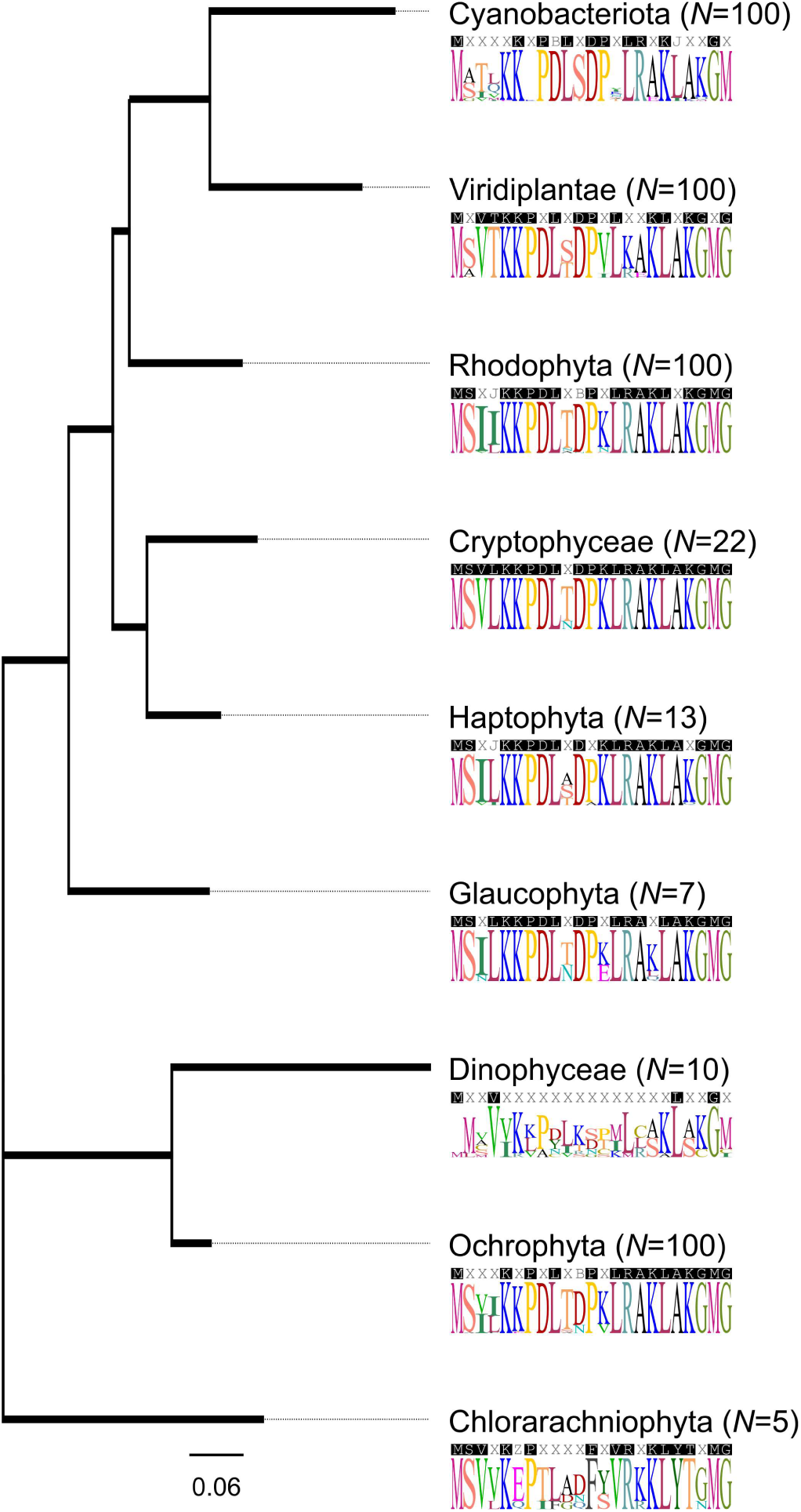
Alignments and phylogenetic tree of PetD highlights T4 in the green lineage. The consensus sequences were used for tree generation and were based on the first 100 BlastP hits (https://blast.ncbi.nlm.nih.gov/) for the *C. reinhardtii* PetD amino acid sequence. Hits were filtered for a Max Score above 100, followed removal of duplicate annotated sequences. The first 23 amino acids are shown.

**Extended Data Table 1. Phosphorylation data, protein fold changes, and primers for *STT7* and *petD* mutant backgrounds.** Available online.

